# A Patient-Derived Cellular Model for Huntington’s Disease Reveals Phenotypes at Clinically Relevant CAG Lengths

**DOI:** 10.1101/291575

**Authors:** Claudia Lin-Kar Hung, Tamara Maiuri, Laura Erin Bowie, Ryan Gotesman, Susie Son, Mina Falcone, James Victor Giordano, Virginia Mattis, Trevor Lau, Vickie Kwan, Vanessa Wheeler, Jonathan Schertzer, Karun Singh, Ray Truant

## Abstract

The huntingtin protein participates in several cellular processes that are disrupted when the polyglutamine tract is expanded beyond a threshold of 37 CAG DNA repeats in Huntington’s disease (HD). Cellular biology approaches to understand these functional disruptions in HD have primarily focused on cell lines with synthetically long CAG length alleles that clinically represent outliers in this disease and a more severe form of HD that lacks age-onset. Patient-derived fibroblasts are limited to a finite number of passages before succumbing to cellular senescence. We used human telomerase reverse transcriptase (hTERT) to immortalize fibroblasts taken from individuals of varying age, sex, disease onset and CAG repeat length, which we have termed TruHD cells. TruHD cells display classic HD phenotypes of altered morphology, size and growth rate, increased sensitivity to oxidative stress, aberrant ADP/ATP ratios and hypophosphorylated huntingtin protein. We additionally observed dysregulated ROS-dependent huntingtin localization to nuclear speckles in HD cells. We report the generation and characterization of a human, clinically relevant cellular model for investigating disease mechanisms in HD at the single cell level, which, unlike transformed cell lines, maintains TP53 function critical for huntingtin transcriptional regulation and genomic integrity.

## INTRODUCTION

Huntington’s disease (HD) is a late-onset, autosomal-dominant neurodegenerative disorder characterized by a triad of motor, cognitive and psychiatric symptoms. The disease is caused by a CAG trinucleotide expansion of >37 repeats in the huntingtin gene, manifesting as polyglutamine-expanded huntingtin protein^1^. The functional implications of this expanded, mutant huntingtin are not fully understood. Much of the existing research on HD cell biology in relevant neuronal cell types has been limited to primary post-mitotic neurons from murine brain tissue or transformed cell lines, which have several limitations, including the use of synthetically long CAG lengths in order to mimic human disease in mice^2–15^. These alleles actually genetically model juvenile or Westphal variant HD, which are not age-onset diseases. The mean clinical CAG allele length is 43 repeats, with even >50 repeats representing statistical outliers^16^. Disease models in Neuro-2A cells, HEK293 or HeLa cells rely on cell transformation to maintain line longevity, but transformation typically involves initiating genomic instability and “shattering” of genomes^17,18^, affecting intra- and inter-laboratory reproducibility.

The pursuit of investigating human cells has driven researchers to culture patient fibroblast cells that can be extracted from a skin biopsy^19–21^. Primary fibroblasts from HD patients possess clinically relevant polyglutamine expansion lengths, making them an attractive model for studying HD human cell biology. Further, they can be reprogrammed into induced pluripotent stem cells (iPSCs)^22^, which can be differentiated into various neuronal cell lineages^23–25^ or directly reprogrammed to medium spiny neurons^26–29^. However, primary fibroblasts are subject to telomere-controlled senescence, which limits the number of passages as telomeres shorten with each cell division^30,31^. Senescent cells display altered gene expression, decreased proliferation and resistance to apoptotic mechanisms^32,33^, hindering long-term use and consistency between trials.

We sought to overcome these limitations by immortalizing patient fibroblasts with human telomerase reverse transcriptase (hTERT). hTERT has been extensively used to immortalize human cell types to study cell biology in a number of diseases^34–39^. Like primary cells, hTERT-immortalized cells mimic *in vivo* tissue phenotypes^40^, with the added benefits of proliferative capacity, karyotypic stability, and inter-experimental reproducibility^40^.

We sought to generate a panel of human cell lines with polyglutamine expansions in the 40-50 CAG range, reflective of those seen in the clinic. We immortalized fibroblasts from 3 individuals and termed the cell lines TruHD cells: control female (TruHD-Q21Q18F), heterozygous male (TruHD-Q43Q17M), and homozygous female (TruHD-Q50Q40F) (Table 1). To validate these cells as HD models, we examined known model and patient phenotypes. Consistent with previous reports, we found that HD cells could be distinguished from wild type based on morphology^7,15^, size^15^, growth rate^20,41,42^, sensitivity to stress^9,43–45^, aberrant ADP/ATP ratios^7,46,47^, hypophosphorylated huntingtin protein^48–50^ and altered ROS-dependent localization to nuclear speckles^51^. We have therefore generated and characterized a panel of clinically relevant cellular models for investigating disease mechanisms in HD.

**Table 1:** TruHD Patient Fibroblast Information

| Primary Cell Name | Cell Line Name | Allele 1 | Allele 2 | Sex | Age at Sampling | Disease Onset Age |
| --- | --- | --- | --- | --- | --- | --- |
| ND30013 | TruHD-Q43Q17M | 43 CAG | 17 CAG | Male | 54 | 50 |
| ND30014 | TruHD-Q21Q18F | 21 CAG | 18 CAG | Female | 52 | - |
| GM04857 | TruHD-Q50Q40F | 50 CAG | 40 CAG | Female | 23 | 28 |

Genome-wide association studies (GWAS) in the HD research field revealed DNA damage and oxidative stress mechanisms as critical modifiers of the age of disease onset in patients^52^. One of the most important regulators of DNA repair, cell stress, and cell death responses is the TP53 protein^53–55^, which also directly regulates huntingtin transcription via a response element in the *HTT* promoter region^56,57^. To date, the most widely used HD cell lines are SV40 large T-antigen-transformed mouse striatal cell lines (ST*Hdh*)^7^, which have become an invaluable tool for studying HD cell biology. However, conditional cell immortalization by transformation requires inhibited TP53 function. Investigation of the role of huntingtin in DNA damage and cell stress pathways may therefore be confounded by TP53 inactivation. In contrast, hTERT immortalization does not alter TP53 function^35,58,59^, making TruHD cells an attractive model for these disease mechanisms in particular. Immortalized fibroblasts provide the added benefits of inter-experimental and inter-laboratory reproducibility, and long-term applications such as generation of stable cell lines and direct conversion to patient-specific human neurons. These cell models are readily available to the HD research community and will be freely distributed by our group.

## RESULTS

### Immortalization of Primary Fibroblasts with hTERT

Primary fibroblasts from various patients were obtained from the Coriell Institute for Medical Research and transduced with TERT Human Lentifect™ Purified Lentiviral Particles as described in methods. Three immortalized cell lines were generated from patients with varying CAG repeat lengths and disease onset age (Table 1) as representatives of control (TruHD-Q21Q18F), heterozygous HD (TruHD-Q43Q17M) and homozygous HD (TruHD-Q50Q40F). We standardized the line naming to a format compatible with digital file annotation, defining the CAG length of each *HTT* allele and the sex of the donor.

To verify that cells were successfully overexpressing hTERT, RNA levels in primary cells and TruHD cells were compared by quantitative PCR (qPCR), showing detectable hTERT mRNA levels in TruHD cells compared to primary cells normalized to commercially available hTERT-immortalized retinal pigment epithelial (RPE1) cells (Figure 1A). To ensure that the increased hTERT expression was associated with increased hTERT catalytic activity, telomerase activity was tested in TruHD-Q21Q18F and TruHD-Q43Q17M cells using a telomeric repeat amplification protocol (TRAP) assay. As shown in Figure 1B, multiple amplification products resulting from active hTERT were observed in TruHD cells, but not primary cells, indicating that the transduced hTERT is catalytically active in TruHD cells.

**Figure 1:**
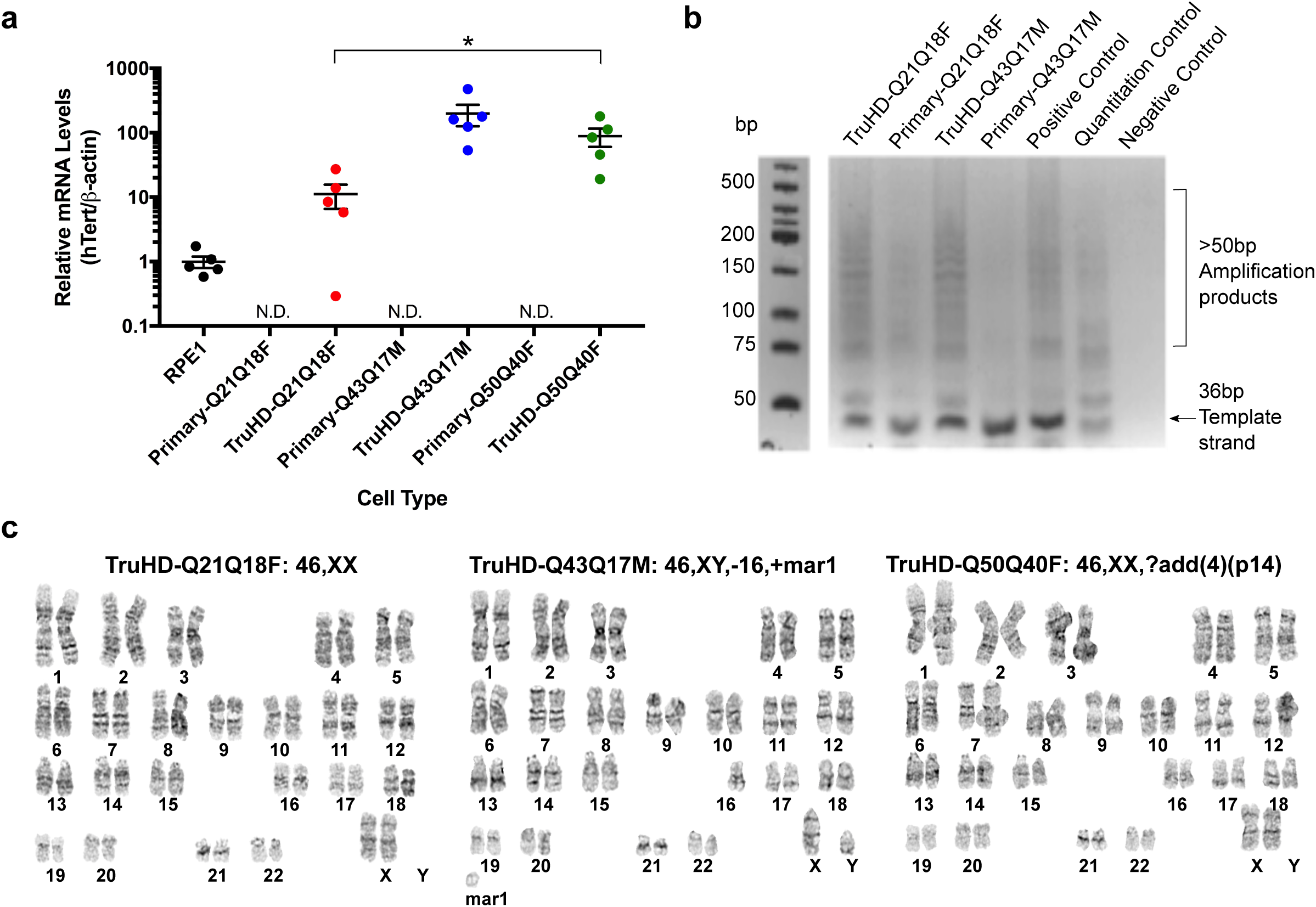
Generation of TruHD immortalized cell lines. (A) hTERT mRNA levels normalized to beta-actin (β-actin) mRNA levels in RPE1 cells (positive control), primary cells and TruHD cells. hTERT levels in primary cells were not detectable (N.D.). n=5. Error bars represent S.E.M. *p=0.0369 comparing TruHD-Q21Q18F, TruHD-Q43Q17M and TruHD-Q50Q40F by one-way ANOVA. (B) Telomeric repeat amplification product (TRAP) assay. Amplification products run on 10% TBE gel after telomere extension reaction, showing telomeric repeats >50 bp in increments of 6 bp. Template strand is 36 bp. Negative control contains no Taq polymerase or template strand. (C) Representative karyotypes of TruHD-Q21Q18F, TruHD-Q43Q17M and TruHD-Q50Q40F cells. “mar” denotes marker chromosomes, “+” are additional chromosomes and “?add(4)(p14)” denotes additional patterns observed on chromosome 4 at band p14. Results from full karyotype shown in Table 2.

Unlike immortalization by transformation, hTERT immortalization maintains karyotypic stability in normal, human diploid cells^35,38,58^. Chromosomal instability leading to polyploidy and aneuploidy can affect gene expression and cell viability, which is a hallmark of transformed cancer cells^60,61^. To confirm karyotypic stability in TruHD cells after 25+ passages, we compared the karyotypes of ST*Hdh* cells and TruHD cells. Large chromosomal abnormalities were detected in transformed HD mouse striatal derived cells (ST*Hdh*^Q111/Q111^) cells (Supplemental Figure 1A, Table 2), consistent with a recently published study^15^. No chromosomal changes were recorded for control TruHD-Q21Q18F, and minor chromosomal changes were recorded in TruHD-Q43Q17M and TruHD-Q50Q40F cell lines (Figure 1C, Table 2). In TruHD-Q43Q17M cells, the majority of the analyzed cells were missing one chromosome 16, and in TruHD-Q50Q40F cells, most of the cells had an abnormal banding pattern on chromosome 4. These changes should be considered when interpreting results of phenotypic analysis.

**Table 2:**
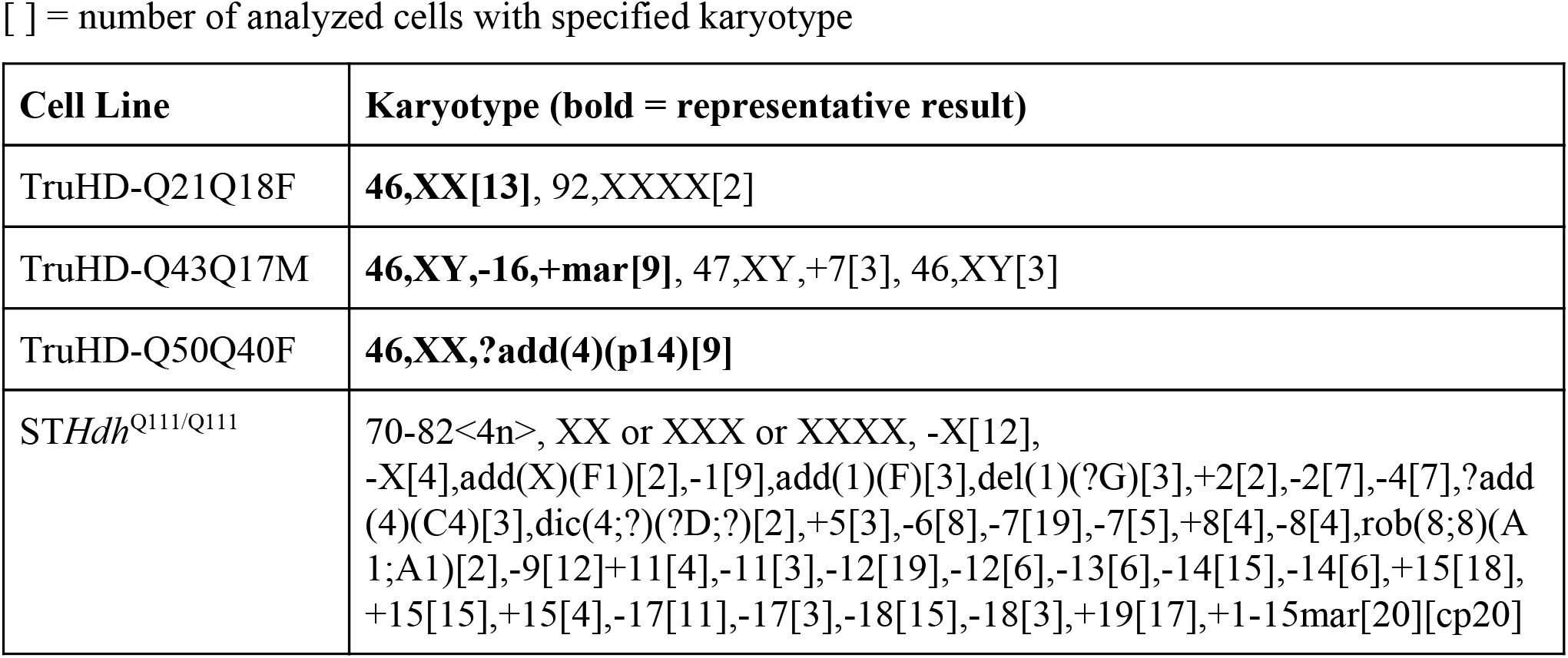
G-band Karyotyping

### Validation of CAG Repeats in TruHD Cell Lines

To verify that the CAG repeats of the fibroblasts matched the clinical information reported after successful immortalization, the CAG repeats were sized using a standardized *HTT* CAG repeat sizing assay^62,63^. The length of each CAG tract was as expected (Table 3). The human huntingtin polyglutamine tract bears an additional CAACAG sequence beyond the pure CAG DNA tract sequence^1^. These two additional codons encoding glutamine residues were not considered in the annotation by the Coriell Institute. Therefore, TruHD-Q21Q18F, for example, only refers to the polyglutamine tract that corresponds to the pure CAG tract, but the full polyglutamine tract lengths are actually Q23Q20. The true polyglutamine lengths corresponding to each TruHD cell line are listed in Table 3.

**Table 3:** Sizing of CAG and CCG repeats in TruHD Fibroblasts

| Cell Line Name | CAGn Repeats | Q lengths (= CAG +2, based on genetic structure of CAGnCAACAG in humans) |
| --- | --- | --- |
| TruHD-Q21Q18F | 21 CAG/18 CAG | 23Q/20Q |
| TruHD-Q43Q17M | 43 CAG/17 CAG | 45Q/19Q |
| TruHD-Q50Q40F | 50 CAG/40 CAG | 52Q/42Q |

### Defining Senescence in TruHD Cell Lines

Primary fibroblasts are typically cultured for approximately 15 passages from the initial skin biopsy before reaching senescence, while the successfully immortalized cells reported here can be passaged beyond 80 passages without reaching senescence (data not shown). Senescent cells show changes in cell growth, morphology and gene expression, which can be detected by an associated beta-galactosidase activity^64,65^. Senescence was not detectable in TruHD immortalized cell lines (Supplemental Figure 1B) under normal culture conditions. We did note, however, that control TruHD-Q21Q18F fibroblasts seeded too sparsely exhibited senescence-associated beta-galactosidase activity (Supplemental Figure 1C) and stopped dividing. Normal, adherent cells in culture undergo contact inhibition, or post-confluence inhibition of cell mitosis^64,66,67^. Essentially, once the cells become too confluent and make contact with nearby cells, they stop dividing and do not grow because of contact inhibition, unlike transformed cell lines. Cells that are left in this state for too long can become senescent and do not recover in culture^68^. Specific protocols for freeze/thaw were therefore considered and explained in detail in the methods section. After 7 days of confluence without media changes, control cells were more susceptible to senescence compared to the HD cell line (Supplemental Figure 1D). Upon karyotypic analysis of TruHD-Q21Q18F cells cultured under these conditions, a small percentage of control cells displayed tetraploidy (Supplemental Figure 1E), a phenomenon which has been reported to be a result of cellular senescence^64,69^. In contrast, tetraploidy did not occur for either of the HD cell lines. Overall, we observed that cells did not senesce after extended passaging (over 80 passages), but control cells did senesce if cultured too sparsely or at high confluence, unlike HD cells which have a more senescence-resistant phenotype. This aspect of these cell lines could provide utility to study huntingtin biology in senescent cells, or cells in transition from mitotic to senescent.

### Cell Morphology, Size, Growth and Viability

Mutant ST*Hdh* cells exhibit altered morphology^7,15,44^. To probe whether TruHD cells could be distinguished by their morphology, nuclei were stained with Hoechst and immunofluorescence was performed with antibodies against phosphorylated huntingtin at serines 13 and 16 (N17-phospho) and beta-tubulin (Figure 2A). Images were analyzed with Phenoripper open software (www.phenoripper.org), which defines textures of the images in a non-supervised manner, and plots vectors of the three most variant textures in unitless 3D space using principal component analysis (PCA). This allows for identification of similarity between the images based on those defined features. Merged images of TruHD-Q21Q18F, TruHD-Q43Q17M and TruHD-Q50Q40F, considering the three parameters Hoechst, N17-phospho, and beta-tubulin, clustered apart from each other on a PCA plot (Figure 2B). The clustering pattern from each individual channel is shown in Supplemental Figure 2A, where beta-tubulin alone showed the greatest separation in the clusters compared to Hoechst and N17-phospho. This unbiased detection of a morphology phenotype between control and HD cells may be attributed to differences in cell size. We therefore compared cell size in control and mutant TruHD cells. Quantification of cell surface area shows that HD cells are smaller than control cells (Figure 2C). Therefore, consistent with numerous independent previous reports, huntingtin may be involved in cytoskeletal regulation^15,70–72^.

**Figure 2:**
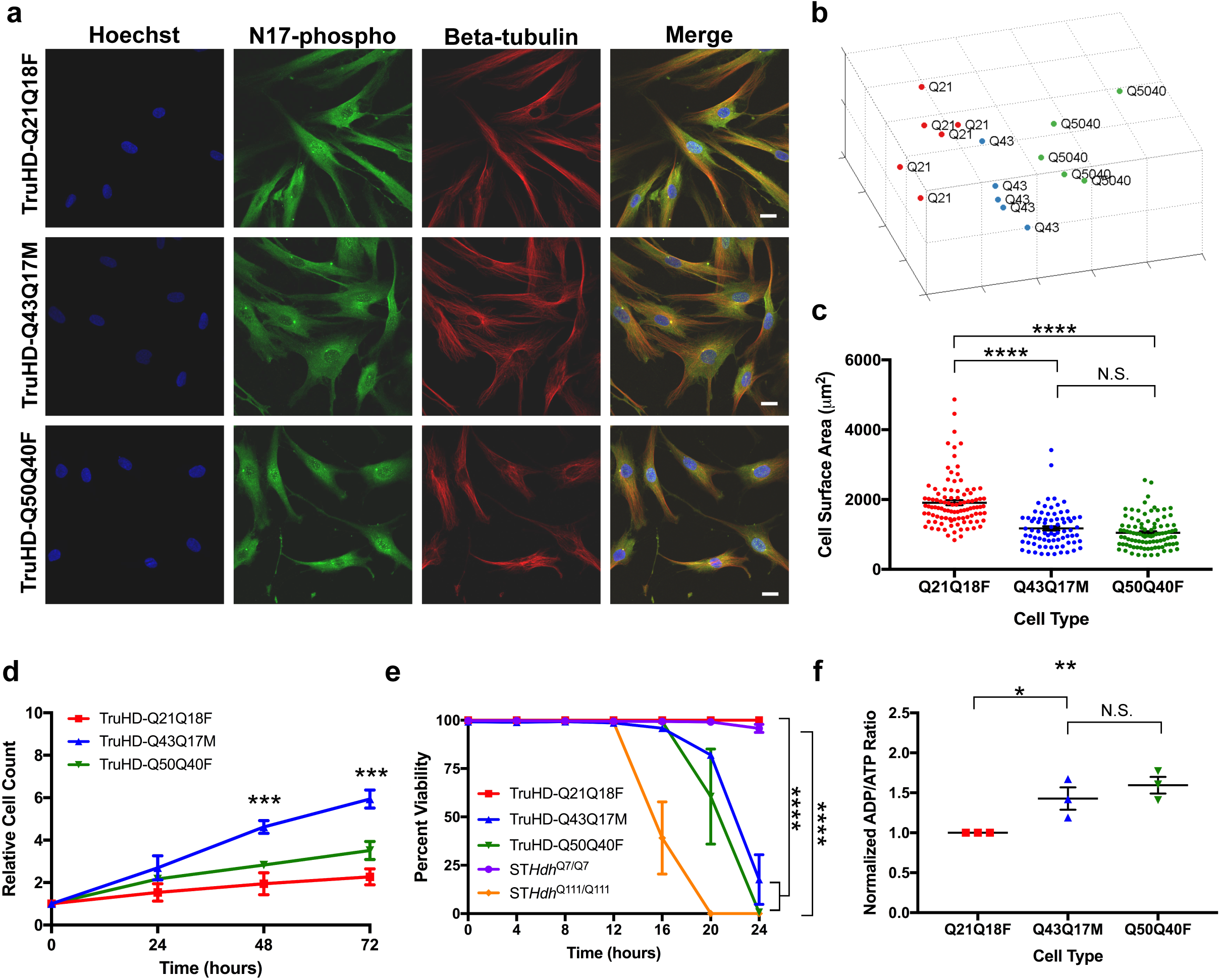
TruHD cell properties. (A) Immunofluorescence images of TruHD-Q21Q18F, TruHD-Q43Q17M and TruHD-Q50Q40F. Scale bar = 10μm. (B) PCA plot of images sorted with Phenoripper. (C) Cell surface area comparison in TruHD cells. n=3, N>200. Error bars represent S.E.M. *p <0.0001. (D) Relative cell count measured every 24 hours. n=3, N>200. Error bars represent S.E.M. ***p=0.0003 at 48 hours, ***p=0.0001 at 72 hours by one-way ANOVA. (E) Percent cell viability of TruHD cells compared to ST*Hdh* cells. n=3, N>200. Error bars represent S.E.M. At 24 hours, ****p<0.0001 for ST*Hdh*^Q7/Q7^ vs ST*Hdh*^Q111/Q111^ by two-ANOVA and ****p<0.0001 for TruHD-Q21Q18F vs TruHD-Q43Q17M and TruHD-Q50Q40F by two-way ANOVA. (F) Normalized ADP/ATP ratio in TruHD cells at ~75% confluency 24 hours after seeding. n=3, N>200. Error bars represent S.E.M. *p=0.0371 and **p=0.0048.

Re-entry of post-mitotic neuronal cells into the cell cycle is a potential mechanism in the neurodegenerative process (see review^73^). A consistent observation when culturing TruHD fibroblasts is that the mutant fibroblasts divide more rapidly, as seen with primary HD fibroblasts^20^. Monitoring proliferation of TruHD cells over 72 hours showed that TruHD-Q43Q17M cells double after ~20 hours, TruHD-Q50Q40F cells double after ~24 hours, while TruHD-Q21Q18F double after ~72 hours (Figure 2D). These observations are consistent with reports implicating huntingtin in cell cycle regulation and DNA damage repair mechanisms^45,72,74^ and that these functions are aberrant in HD cells.

Susceptibility to various types of cell stress is a well documented phenomenon in HD cellular models^43,49,70,75^. We have previously shown that huntingtin responds to oxidizing agents by becoming phosphorylated, translocating from the ER to the nucleus and interacting with chromatin^45,51^. We therefore tested TruHD cell viability upon oxidative stress with potassium bromate (KBrO_3_) treatment over 24 hours. As shown in Figure 2E, both HD cell lines were most susceptible to cell death compared to the control line. Dose-dependent response of treatments in each individual TruHD cell line can be found in Supplemental Figure 2B.

Besides response to cell stress, another well documented HD phenotype is an energy deficit as measured by ADP/ATP ratio^7,44,46^. The detection of ADP/ATP ratio in TruHD cells demonstrated an energy deficit in HD cell lines compared to the wild type cell line (Figure 2F) similar to the trend in ST*Hdh* cells (Supplemental Figure 2C), which is consistent with previously reported studies in HD models and synthetic allele lengths^7,47^.

These phenotypes described in TruHD cells such as cell morphology, size, growth rate, susceptibility to oxidative stress and energy levels, demonstrate their utility as a cellular model with clinically relevant CAG allele lengths as phenotypes in mutant TruHD cells were all consistent with previous HD cellular models.

### Total Huntingtin and Phosphorylated Huntingtin Protein Levels

Since the cells were taken from various patients and are not isogenic, the amount of total huntingtin was quantified. Validated antibodies to different epitopes of full-length huntingtin were used (EPR5526 and mAb2166), showing a minor decrease in total huntingtin levels in TruHD-Q50Q40F compared to TruHD-Q21Q18F and TruHD-Q43Q17M (Figure 3A, B; full immunoblots Supplemental Figure 4A, B).

**Figure 3:**
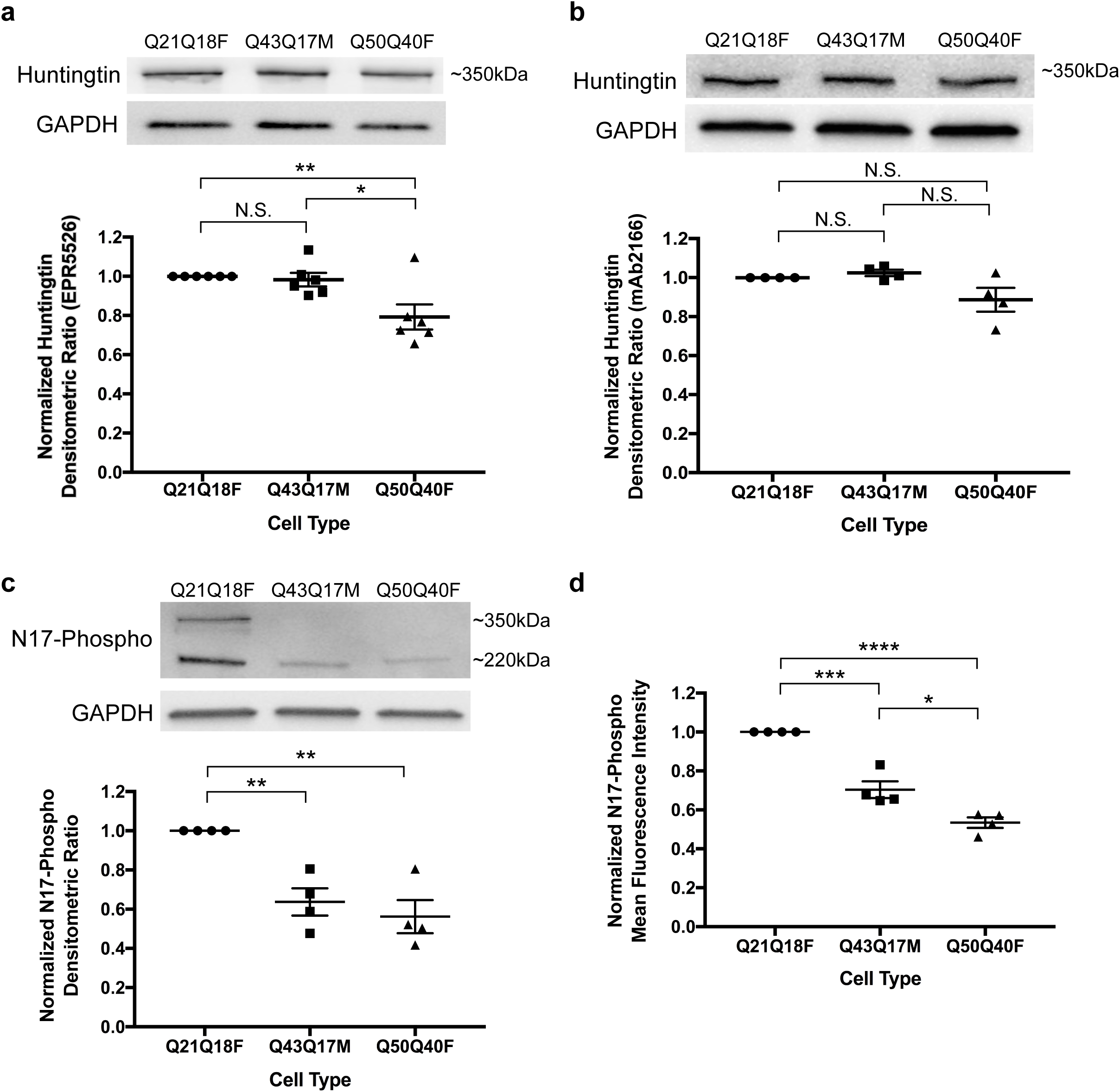
Huntingtin protein levels in TruHD cells. (A) Densitometric analysis of total huntingtin levels using western blot with EPR5526 antibody. Immunoblots were cut horizontally at 75 kDa marker so that GAPDH loading control was probed for separately. Normalized to control TruHD-Q21Q18F cells. n=6. Error bars represent S.E.M. **p=0.0087 and *p=0.0254 by unpaired t-test. (B) Densitometric analysis of total huntingtin levels using western blot with mAb2166 antibody. Immunoblots were cut horizontally at 75 kDa marker so that GAPDH loading control was probed for separately. Normalized to control TruHD-Q21Q18F cells. n=4. Error bars represent S.E.M. (C) Densitometric analysis of N17-phospho levels using western blot. Immunoblots were cut horizontally at 75 kDa marker so that GAPDH loading control was probed for separately. Normalized to control TruHD-Q21Q18F cells. n=4. Error bars represent S.E.M. **p=0.0020 by unpaired t-test. (D) Mean fluorescence intensity analysis of N17-phospho using flow cytometry. Normalized to control TruHD-Q21Q18F. n=4. Error bars represent S.E.M. ***p=0.0005, *p=0.0159 and ****p<0.0001 by unpaired t-test.

Phosphorylation is a protective post-translational modification in HD cells^48–50^. Restoration of N17 phosphorylation is a therapeutic target in HD that has been explored by our lab and others because mutant, polyglutamine-expanded huntingtin is hypophosphorylated at serines 13 and 16 (S13p and S16p respectively)^48–50^. Immunoblotting performed with a validated antibody against both serines (α-N17-phospho) (Supplemental Figure 3) showed decreased levels of N17 phosphorylation in mutant TruHD fibroblast cell lines compared to wild type (Figure 3C; full immunoblot Supplemental Figure 4C). Two distinct molecular weight bands are seen with the antibody, at ~350kDa and ~220kDa. Degradation products are often reported with other huntingtin antibodies as well, due to the large size of the protein and the rigorous processing steps of immunoblotting^76,77^. Both bands were considered in quantifications, and this confirms hypophosphorylation of mutant huntingtin in a human HD. These results were verified by measuring whole-cell mean fluorescence intensity by flow cytometry (Figure 3D). Therefore, N17-phospho levels vary, but total huntingtin levels are invariant. This phenotype is consistent with previous reports^48–50^ and further validates N17-phosphorylation restoration as a target for HD therapeutic development.

### Huntingtin Stress Response in Human Patient Fibroblasts

Huntingtin is a stress response protein and is involved in the unfolded protein response (UPR), DNA damage repair, oxidative stress and endoplasmic reticulum (ER) stress pathways^43,45,51,78,79^. Previous studies from our lab show that huntingtin is bound to the ER membrane in steady state conditions and is released under conditions of stress, particularly ROS stress^43,49^. Once soluble, huntingtin is phosphorylated at serines 13 and 16 (S13,S16), translocates to the nucleus and localizes to nuclear puncta^49,51,80^. Using super-resolution structured-illumination microscopy (SR-SIM), we have now identified that these previously reported nuclear puncta are SC35 positive nuclear speckles: dynamic RNA/protein structures that are rich in mRNA splice factors that are important in cell stress responses^81–83^ (Figure 4A).

**Figure 4:**
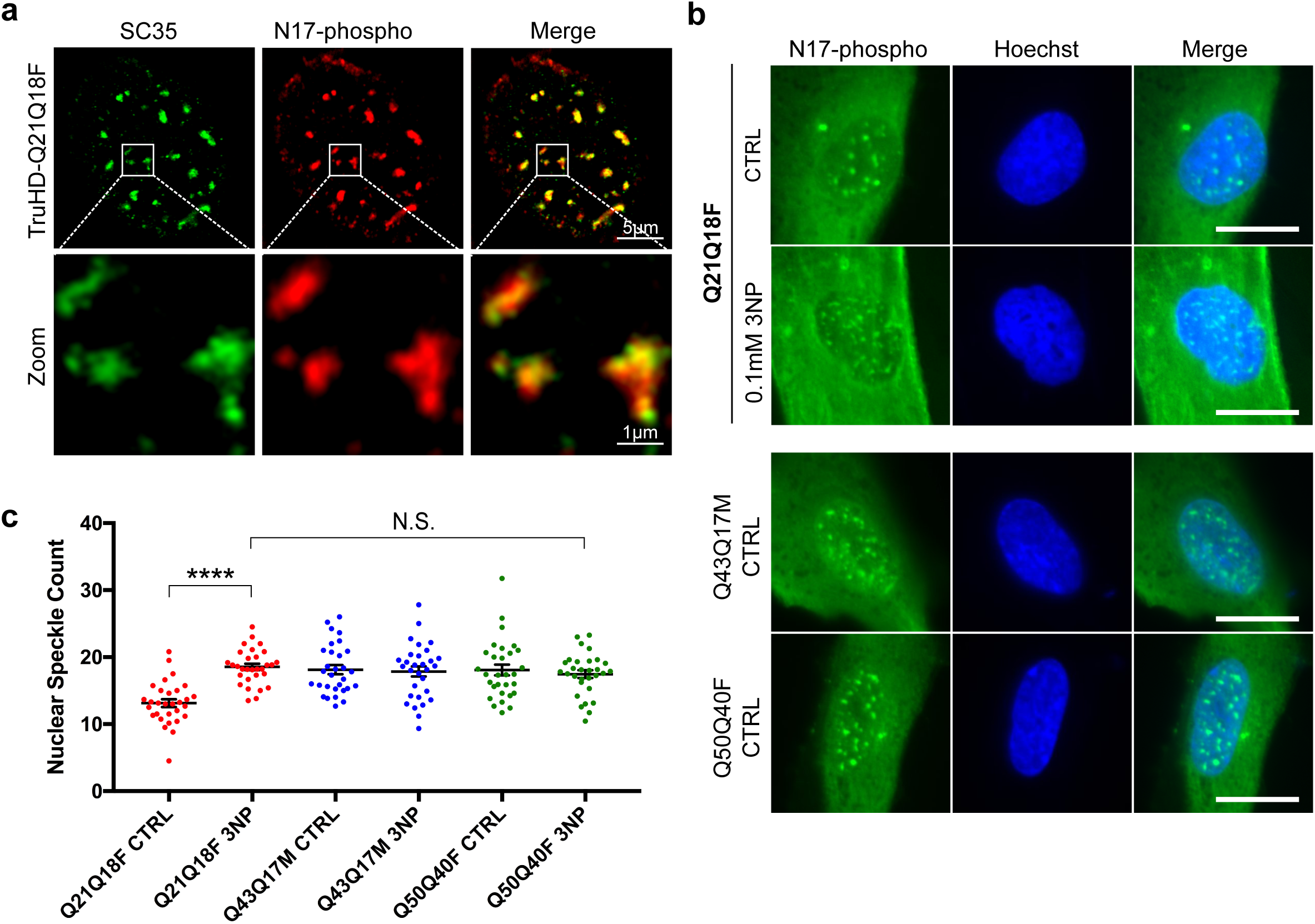
Huntingtin localizes to nuclear speckles in response to stress. (A) N17-phospho localizes to SC35+ nuclear speckles. (B) Nuclear speckles are increased in TruHD-Q21Q18F control cells upon treatment with 0.1mM 3NP, but not for TruHD-Q43Q17M or TruHD-Q50Q40F. Scale bar =10μm. (C) Quantification of nuclear speckles. n=3, N=180. Error bars represent S.E.M. ****p<0.0001 by unpaired t-test. Comparison of TruHD-Q21Q18F 3NP, TruHD-Q43Q17M CTRL, TruHD-Q43Q17M 3NP, TruHD-Q50Q40F CTRL and TruHD-Q50Q40F 3NP by one-way ANOVA shows no significant difference (p=0.8475).

We previously reported that stress-dependent phosphorylation of S13 and S16 promotes huntingtin localization to nuclear speckles in hTERT-immortalized human retinal pigment epithelial cells (RPE1)^51^. We therefore tested this phenomenon in TruHD cells. Cells were treated with 0.1mM 3-nitroproprionic acid (3NP), a mitochondrial complex II inhibitor, for 1 hour to induce oxidative stress. We observed a significant increase in the number of nuclear speckles in TruHD-Q21Q18F cells (Figure 4B, C). However, in both the mutant TruHD-Q43Q17M and TruHD-Q50Q40F lines, there was no significant difference in the number of nuclear speckles between treated and untreated conditions. The number of nuclear speckles in both heterozygote and homozygote HD cells were similar to that of TruHD-Q21Q18F in the presence of 3NP (N.S., p=0.8475), suggesting that in HD cells cells are under a chronic stress load.

Characterization of several HD phenotypes, combined with the establishment of methods to easily detect these disease-relevant phenotypes, in TruHD cells demonstrate their utility as a cellular model and will hopefully facilitate further investigation into pathological mechanisms.

## DISCUSSION

Patient fibroblasts have been used previously by us and others^19,41,42,45^ to study HD cell biology, but currently there are no defined cell lines that are used consistently between projects. After generation of hTERT-immortalized TruHD cells, we focused on defining characteristics of these cells in order to facilitate their use. Our wild type line (TruHD-Q21Q18F), heterozygous HD line (TruHD-Q43Q17M) and homozygous HD line (TruHD-Q50Q40F) were chosen for this study as their CAG repeat lengths were most representative of annotated lengths, but also because they cultured well and the observed phenotypes were consistent throughout the study. A homozygote TruHD-Q50Q40F was chosen, because although a rare clinical example, this line could have utility as both alleles of huntingtin are mutant expanded, and thus can help resolve data in heterozygote lines, where mutant huntingtin phenotypes could be confounded by the presence of the normal allele.

We observed HD phenotypes in both heterozygous TruHD-Q43Q17M and homozygous TruHD-Q50Q40F cells, at clinically relevant CAG repeat lengths. Typical HD cell phenotypes include reduced cell size^15^, decreased cell viability upon cellular stress^43,49,84^, altered cell proliferation^15^, decreased ADP/ATP ratio^7^ and hypophosphorylation of huntingtin N17 at serines 13 and 16^48–50^. Here, we have also demonstrated that HD cells showed altered susceptibility to cellular senescence and deficient response to oxidative stress as seen by SC35 nuclear speckle counts. Additionally, TruHD cells can be distinguished in an unbiased manner using non-supervised image texture analysis and principal component analysis via software such as Phenoripper. Phenoripper can be used as a readout for future high-content drug screening assays. Improvement in inter-experimental and inter-laboratory reproducibility have also been observed and may be beneficial for long-term applications such as generation of stable cell lines and cellular reprogramming to generate patient-specific neurons that maintain age-associated signatures^26,28,29^.

Recent developments in understanding cell biology throughout the course of HD progression highlight the need for improved methods for disease modelling. DNA damage repair pathways have been implicated as the predominant modifiers of HD pathogenesis^52^. Previous observations in our lab show that huntingtin can sense oxidative stress and that huntingtin is involved in the DNA damage response^45^. These processes require TP53 which an important transcription factor that integrates various cellular stress signals and is widely considered the master regulator of genomic integrity due to its roles in DNA damage sensing, cell cycle checkpoint control and apoptotic regulation (see reviews^85–87^).

The choice of model system for studying certain aspects of cell biology is therefore critical. Historically, synthetically long CAG repeat alleles of *HTT* have been used in cell models because of the apparent lack of obvious phenotypes of clinical HD alleles in animal models. Cell biology research in HD has been primarily focused on neurons from HD mouse models and easily accessible cell lines, each with their own restrictions and limitations. The main limitation of cells taken from mouse models are the synthetically long polyglutamine tracts used in order to mimic a late-onset human disease within the lifespan of a mouse. In these models, it can be overlooked that the majority of patients have CAG repeats between 40-50, and that these patients have varying age onset that is not attributed to just the number of repeats. Additionally, some models are transgenic, and thus do not have an accurate gene dosage, while others express huntingtin at super-physiological levels, confounding data with incorrect protein stoichiometry, which can be a concern for a scaffolding protein. This is the first characterized human HD immortalized cell line model and can therefore be used to test therapeutic reagents that are designed specifically for human cells and will be a tool for the HD research community.

## METHODS

### Cell Culture and Generation of hTERT-Immortalized Fibroblasts

Patient fibroblasts were purchased from the Coriell Institute from the NINDS repository. HD patient fibroblasts (ND30013, GM04857) and control patient fibroblasts (ND30014) were obtained. Cells were cultured in Minimum Essential Media (MEM, Gibco #10370) with 15% fetal bovine serum (FBS, Gibco) and 1X GlutaMAX (Gibco #35050). Cells were infected with 1 x 10^6^ TERT Human Lentifect™ Purified Lentiviral Particles (GeneCopoeia, LPP-Q0450-Lv05-200-S). To aid in infection, 10 μg/mL polybrene was added. After 8 hours, cells were infected again and left for 24 hours. Media was changed and cells were left for an additional 48 hours. Successfully transduced cells were selected in media with 1 μg/mL puromycin. Cells were grown at 37°C with 5% CO_2_.

ST*Hdh* cells (a kind gift from Dr. Marcy Macdonald) were cultured in Dulbecco’s Modified Eagle Medium (DMEM, Gibco #11995) with 10% FBS. Cells were grown at 33°C with 5% CO_2_. RPE1 cells (ATCC) were cultured in 1:1 Dulbecco’s Modified Eagle Medium/Nutrient Mixture F-12 (DMEM/F12, Gibco #11330) with 10% FBS and 0.01% hygromycin. Cells were grown at 37°C with 5% CO_2_.

### Quantitative PCR to Measure hTERT mRNA Levels

Primary, TruHD and RPE1 cells were grown to ~85% confluence. Total RNA was obtained from frozen cell pellets lysed in 1 mL of Trizol (Thermo Fisher Scientific) per ~1 x 10^6^ cells, followed by phenol-chloroform extraction. RNA was treated with DNase I (Thermo Fisher Scientific) and cDNA was prepared using 1000 ng total RNA and SuperScript III Reverse Transcriptase (Thermo Fisher Scientific). Transcript expression was measured using TaqMan Assays with AmpliTaq Gold DNA polymerase (Thermo Fisher Scientific), and *hTERT* (Hs00972650_m1) was compared to *ACTB* (Hs01060665_g1) housekeeping gene using the ΔΔ*C_T_* method.

### Telomeric Repeat Amplification Protocol

Primary fibroblasts and corresponding TruHD cells were grown to ~85% confluence. A two-step PCR method (TRAPeze^®^ Telomerase Detection Kit S7700, Millipore) was used to evaluate hTERT catalytic activity. Briefly, in the first step, telomerases from lysed cells add telomeric repeats (AG followed by repetitive GGTTAG sequences) on the 3’ end of a substrate oligonucleotide (TS). In the second step, the extended products are amplified by PCR using a primer specific for the beginning of TS, and a reverse primer specific for the end of the repeats. The lowest amplification product should be 50 bp, and increases by 6 bp-long repeat increments are visualized, on a 10% TBE polyacrylamide gel.

### Cryogenic Storage of TruHD Cells

For freezing one vial of TruHD cells, a plate was grown to 90% confluence (~1 x 10^5^/mL) on a 10 cm dish and was split in half. The next day, two ~60% confluent plates, that were still in growth phase, were trypsinized and combined. Cells were centrifuged at 1500 rpm and pellets were resuspended in culture media with 1 mL of 5% DMSO. Vials were put into a slow-freeze unit in the -80°C to ensure optimal cell preservation. After 24-48 hours, vials were moved to a -150°C for long-term storage.

Vials were thawed slowly at 37°C for around 2-5 minutes. Using a 10 mL pipette, the 1 mL of cells were moved directly into a 10 cm plate already pre-incubated with media. After 24 hours, media was changed to remove residual DMSO.

### Sizing of CAG Repeat

TruHD cells were grown to ~90% confluence in a 10 cm plate. Cells were scraped and centrifuged at 4°C at 1500 rpm. Genomic DNA was extracted using the PureLink^®^ Genomic DNA kit (ThermoFisher). A fluorescence-based assay was used to size the CAG repeats, based on the originally described assay by Warner et al.^63^ and further described in Keum et al.^62^

### Karyotyping

Karyotyping was performed by The Centre for Applied Genomics, The Hospital for Sick Children, Toronto, Canada. Karyotype analysis via G-banding was performed on cells from two T25 flasks per cell line. When cells reached 80-90% confluence, Karyomax Colcemid^®^ was added to each flask to a final concentration of 0.1 μg/mL (Gibco #15212-012) and incubated in 37°C CO_2_ incubator for 1.5-2 hours (for TruHD-Q43Q17M and TruHD-Q50Q40F) and 3-4 hours for TruHD-Q21Q18F. Cells were then collected and suspended in 6 mL of 0.075 M KCl, and incubated at 37°C for 20 minutes. Eight drops of Carnoy’s Fixative (methanol/acetic acid, 3:1) was added and mixed together. The cells were centrifuged at 1000 rpm for 10 minutes at room temperature and cell pellets were collected. After three rounds of fixations (add 8 mL fixative and centrifuge at 1000 rpm for 10 minutes), cells were resuspended in 0.5-1 mL of fixative and cells from each suspension were dispensed onto glass slides and baked at 90°C for 1.5 hours. Routine G-banding analysis was then carried out. Approximately 15-20 metaphases per cell line were examined.

### Senescence-Associated Beta-Galactosidase Activity Assay

TruHD and primary fibroblasts were seeded into a 6 well plate. Primary fibroblasts reached ~20 passages before analysis and TruHD cells reached ~50 passages. When cells reached ~90% confluency, media was aspirated and cells were washed 1X with PBS and experiments were carried out using Senescence Detection Kit (Abcam, ab65351) manufacturer instructions.

### Phenoripper and Cell Surface Area Measurement

Immunofluorescence was performed with Hoechst 33342 (ThermoFisher), beta-tubulin antibody (E7, DSHB, 1:250 dilution in 2% FBS in PBS with 0.02% Tween) and N17-phospho antibody (NEP, see section on antibody validation) were analyzed using Phenoripper. Five images per trial of TruHD fibroblasts acquired on a Nikon TiEclipse inverted epifluorescent widefield using a 20X objective (NA=0.75) and Spectra X LED lamp (Lumencor) capture using an Orca-Flash 4.0 CMOS camera (Hamamatsu). Cell surface area was calculated with ImageJ. Cells were thresholded to remove background to identify whole-cell region of interest and area of each cell was measured and plotted.

### Cell Counting

Cells were seeded into a 24 well plate (10^5^/mL). After 24 hours, nuclei were stained for 15 minutes with NucBlue Live ReadyProbes Reagent (1 drop/mL of media) (ThermoFisher). Cells were imaged using the Nikon TiEclipse inverted widefield epifluorescent microscope, and using an automated round object detector in NIS Elements Advanced Research 4.30.02v software (Nikon), cell nuclei were counted. This was repeated repeat plates that were left for 48 and 72 hours for all cell lines.

### ADP/ATP Ratio Assay

ADP/ATP Ratio Assay Kit (Sigma MAK135) was used according to the protocol, except the first step to seed TruHD cells directly into a 96 well plate which the rest of the assay is performed on. If seeded directly into a 96 well plate, there is not enough room to grow the suggested 10^4^ fibroblast cells because of their large size. Therefore, TruHD cells were seeded into a 24 well plate, for no more than 48 hours, to ~80% confluency. After reaching confluency, the cells are lysed with the working reagent (provided in the kit). After lysing, cells from the 24 well plate were moved directly into a 96 well plate to continue the rest of the assay according to the protocol. For ST*Hdh* cells, assay was performed according to the protocol with no changes.

### Cell Viability Assay

Cells were seeded into a 96 well plate. After 24 hours, cells were stained for 15 minutes with NucBlue Live ReadyProbes Reagent (1 drop/mL of media) (ThermoFisher) in Hank’s Balanced Salt Solution (HBSS) (Gibco). Cells were washed with once with HBSS and treated with 100 μL of potassium Bromate (KBrO_3_) (Millipore) at concentrations of 0, 1, 10, 100 and 200 mM in HBSS with NucGreen Dead 488 ReadyProbes Reagent (1 drop/mL of media) (ThermoFisher).

The 96 well plate was imaged immediately over the course of 24 hours every 20 minutes at 37°C using a Nikon TiEclipse inverted A1 confocal microscope equipped with a 20X objective (NA= 0.75) and driven by NIS Elements AR 4.30.02v 64-bit acquisition software (Nikon). Cells were imaged simultaneously in the FITC (NucGreen) and DAPI (NucBlue) channels. A cell was defined as undergoing cell death when 50% or more of the nucleus, as defined by NucBlue-positive pixels, was overlapped by NucGreen-positive pixels. Cell death was multiplied by 100% and subtracted from 100 to calculate % Viability. Images were analyzed using Python.

### Immunoblotting

TruHD cells were grown to ~90% confluence. Cells were scraped and centrifuged at 4°C at 1500 rpm. Cell pellets were lysed in radioimmunoprecipitation assay (RIPA) buffer with 10% phosphatase (Roche) and 10% protease inhibitors (Roche) for 12 minutes on ice and centrifuged at 10 000 x g at 4°C for 12 minutes. Supernatant was collected and 40 µg of protein was loaded into a pre-cast 4-20% polyacrylamide gradient gel (Biorad). Proteins were separated by SDS-PAGE and electroblotted onto 0.45 µm poly-vinyl difluoride (PVDF) membrane (EMD Millipore).

Blots were blocked with 5% non-fat dry milk in TBS-T (50 mM Tris-HCl, pH 7.5, 150 mM NaCl, 0.1% Tween-20) for 1 hour at room temperature. Blots were cut horizontally at 75kDa marker to probe for huntingtin (~350kDa) or GAPDH (loading control, ~37kDa) separately. Blots were then incubated with primary N17-phospho antibody (1:1250), EPR5526 (1:2500, Abcam ab106115), mAb2166 (1:2500, Millipore) or GAPDH (1:10 000, Abcam ab8425) overnight at 4°C. Blots were washed 3 times for 10 minutes with TBS-T and then incubated with anti-rabbit or anti-mouse HRP secondary (1:50 000, Abcam) for 45 minutes at room temperature. Finally, blots were washed 3 times for 10 minutes with TBS-T and visualized with enhanced chemiluminescent HRP substrate (EMD Millipore) on a MicroChemi system (DNR Bio-imaging Systems). Huntingtin bands were quantified using NIH ImageJ and normalized to the GAPDH loading control.

### Flow Cytometry

TruHD cells were grown to ~90% confluence. Cells were scraped and centrifuged at 4°C at 1500 rpm. Cells (~10^6^) were resuspended and fixed in ice-cold methanol for 12 minutes, inverting every 4 minutes. Cells were centrifuged at 4°C at 10 000 x g for 5 minutes followed by 2 washes in flow buffer (PBS with 2.5 mM EDTA and 0.5% BSA), and blocked in flow buffer with 2% FBS for 1 hour at room temperature. Cells were incubated overnight in AlexaFluor488-conjugated N17-phospho antibody, diluted 1:15 in flow buffer with 0.02% Tween-20, rotating at 4°C. Cells were washed twice and resuspended in flow buffer.

### 3NP Treatment and Nuclear Speckle Count

TruHD cells were treated with 0.1 mM 3-nitroproprionic acid (3NP) for 1 hour at 37°C, then fixed and permeabilized with ice-cold methanol for 12 min. Cells were washed in PBS and blocked in antibody solution (2% FBS, 0.1% (v/v) Triton X-100 in 1X TBS) at room temperature for 10 minutes. AlexaFluor488-conjugated N17-phospho antibody was diluted 1:15 in antibody solution and incubated overnight at 4°C. Cells were washed in PBS and stained with Hoechst 33342 dye for 12 minutes at room temperature and left in PBS prior to imaging.

Cells were imaged using Nikon TiEclipse inverted widefield epifluorescent microscope using a Plan Apo 60X (NA=1.4) oil objective and Spectra X LED lamp (Lumencor) captured on an Orca-Flash 4.0 CMOS camera (Hamamatsu). A z-stack was obtained for each image and displayed as a maximum projection prior to image analysis. Image acquisition was completed using the NIS-Elements Advanced Research 4.30.02v 64-bit acquisition software (Nikon). Nuclear speckles were quantified in over 200 cells over 3 trials using an open source speckle counting pipeline in CellProfiler (www.cellprofiler.org).

### Dot Blot Assay for Antibody Validation

Varying concentrations (from 25-1000 ng) of different synthetic N17 peptides (N17, N17S13p, N17S16p, and N17S13pS16p) were spotted onto a nitrocellulose membrane (Pall Life Sciences) and allowed to dry at room temperature for 45 minutes. Immunoblotting was carried out as described earlier.

### Immunofluorescence Peptide Competition Assay for Antibody Validation

The N17-phospho antibody (1:250) was incubated, with rotation, with 1000 ng of synthetic N17 peptides (N17, N17S13p, N17S16p, N17S13pS16p, and a control peptide – p53 (371-393); New England Peptides) at room temperature for 1 hour prior to overnight incubation with RPE1 cells fixed with methanol. Cells were washed 3 times with 2% FBS in PBS and then incubated in anti-rabbit AlexaFluor488 secondary antibody (1:500, Molecular Probes) for 45 minutes at room temperature and then washed and left in PBS before imaging using a Nikon TiEclipse inverted epifluorescent microscope.

### Huntingtin siRNA Knockdown for Antibody Validation

Endogenous huntingtin knockdown was established with huntingtin siRNA (Santa Cruz, sc35617) in RPE1 cells. siRNA was transfected using Lipofectamine RNAiMax (Invitrogen) according to manufacturer instructions. Control dishes were transfected with scrambled siRNA. Protein was extracted using as described earlier, and 60 µg protein was loaded. Immunoblotting was carried out as described earlier.

### Statistics

Data with a normal distribution were analyzed by unpaired t-test unless otherwise stated. Error bars represent standard error of the mean (S.E.M.).

### Data Availability

The raw datasets generated and analyzed for this study are available from the corresponding author on reasonable request.

## ACKNOWLEDGEMENTS

The authors wish to thank Raymond Wong of The Centre for Applied Genomics, The Hospital for Sick Children, Toronto, Canada for assistance with G-banding karyotype analysis and Alina Lelic of the Human Immune Testing Suite, McMaster Immunology Research Centre, Hamilton, Canada for assistance with flow cytometry experiments. This work was supported by the Canadian Institutes Of Health Research, grant MOP-119391, the Krembil Foundation and the Huntington Society of Canada.

## AUTHOR CONTRIBUTIONS

C.L.H. and R.T. created experiments. C.L.H. wrote manuscript and generated figures. T.M. and R.T. helped with editing. M.F. provided technical assistance. C.L.H. performed all experiments except for the following: V.M., V.K. and K.S. transduced patient cell lines for immortalization. J.V.G. and V.W. performed CAG repeat sizing. T.L. and J.S. performed qPCR. L.E.B performed antibody validation experiments. R.G. performed cell viability experiments. S.S. performed nuclear speckle count. T.M. performed super-resolution imaging of nuclear speckles.

